# Enhancement of network architecture alignment in comparative single-cell studies

**DOI:** 10.1101/2024.08.30.608255

**Authors:** Clemens Schächter, Martin Treppner, Maren Hackenberg, Hanne Raum, Joschka Bödecker, Harald Binder

## Abstract

Animal data can provide meaningful context for human gene expression at the single-cell level. This context can improve cell-type or cell-state detection and clarify how well the animal models human biological processes. To achieve this, we propose a deep learning approach that identifies a unified latent space to map complex patterns between datasets. Specifically, we combine variational autoencoders with a data-level nearest neighbor search to align neural network architectures across species. We visualize commonalities by mapping cell samples into the latent space. The aligned latent representation facilitates information transfer in applications of liver, white adipose tissue, and glioblastoma cells from various animal models. We also identify genes that exhibit systematic differences and commonalities between species. The results are robust for small datasets and with large differences in the observed gene sets. Thus, we reliably uncover and exploit similarities between species to provide context for human single-cell data.

## 2 Main

Model organisms are crucial in advancing biomedical research by offering advantages like easy genetic manipulation and access to datasets from a variety of experimental contexts [1]. As a popular choice, mouse models have significantly contributed to the study of human diseases [2], including diabetes [3], glioblastoma [4], or non-alcoholic fatty liver disease [5]. However, translating experimental findings to humans is challenging due to biological differences between species. Efforts to bridge this evolutionary gap include engineered mouse models that replicate human biology more closely [6]. Yet, the emergence of single-cell RNA sequencing (scRNA-seq) has also opened up opportunities for deep learning approaches to compare experimental findings across species.

Transfer learning techniques established themselves as powerful tools for sharing information between scRNA-seq datasets. These approaches often use encoder-decoder architectures to compress datasets into a low-dimensional manifold. Examples include Cell BLAST [7] and ItClust [8], which annotate and cluster cells based on knowledge transfer from reference datasets.

Architecture surgery techniques adjust network architectures to the characteristics of different datasets. After pre-training, additional neurons are inserted into the encoder and decoder input layers. They correct for unseen batch effects in the new data, while all other weights remain fixed during subsequent training. This approach, pioneered by scArches [9], now spans a diverse set of models [10–12]. Despite the method’s success, two primary challenges remain unaddressed for datasets of different species. (Figure 4.)

First, some genes lack orthologs in other genomes, which requires different interpretations of certain input nodes in their neural network architectures. For instance, 20% of human protein-coding genes and a significant percentage of small and long non-coding RNAs miss one-to-one mouse orthologs [13]. To enable training, architecture surgery-based approaches restrict datasets to orthologous genes or zero-fill missing values. Outside of architecture surgery, some models like SATURN [14] and TACTiCS [15] match genes via protein sequences with transformer-based language models.

The second challenge is that biological similarities between cells do not always translate into similar gene expression patterns, which can vary significantly between species [13]. Therefore, neural networks may struggle to recognize similar cells.

To account for differences between gene sets and expression levels, we introduce scSpecies. Our approach pre-trains a conditional variational autoencoder-based model [16] and fully re-initializes the encoder input layers and the decoder network during fine-tuning. Architecture alignment is guided by a nearest neighbor search performed on homologous genes, which estimates similarity between cells in both datasets. This incentivizes our model to map biologically related cells into similar regions of the latent space. The neighbor search requires only a small subset of observed genes to be homologs, while all remaining genes can have no relationship at all. Moreover, scSpecies enables nuanced comparisons of gene expression profiles by generating gene expression values for both species from a single latent variable.

We tested our method on data from various species and organs, including liver cells [17], white adipose tissue cells [18], and glioblastoma immune response cells [19, 20]. Our results demonstrate that scSpecies effectively aligns network architectures and latent representations. We improve upon cell label transfer from the initial nearest neighbor search and existing architecture surgery approaches when measured in terms of accuracy and multiple clustering metrics.

## 3 Results

We present scSpecies, a tool for researchers who wish to use one scRNA-seq dataset as context for another from a different species. In the following, the dataset of the model organism is referred to as ‘context dataset’ and the dataset of the target organism as ‘target dataset’. scSpecies aligns context scRNA-seq datasets with human target data, enabling the analysis of similarities and differences between the datasets.

Next to context and target datasets, the model requires a sequence containing indices of homologous genes, indicator variables for batch effects, and cell type labels for the context dataset.

The proposed workflow (Figure 1) aligns the network architectures of two single-cell variational inference (scVI) [21] models in a pre-training strategy. In scVI, encoder neural networks map gene expression vectors into a compressed latent space separating cells by biological features. Conversely, a decoder maps from this low-dimensional representation onto parameters of a negative binomial distribution to (re-)generate gene expression data.

**Figure 1:**
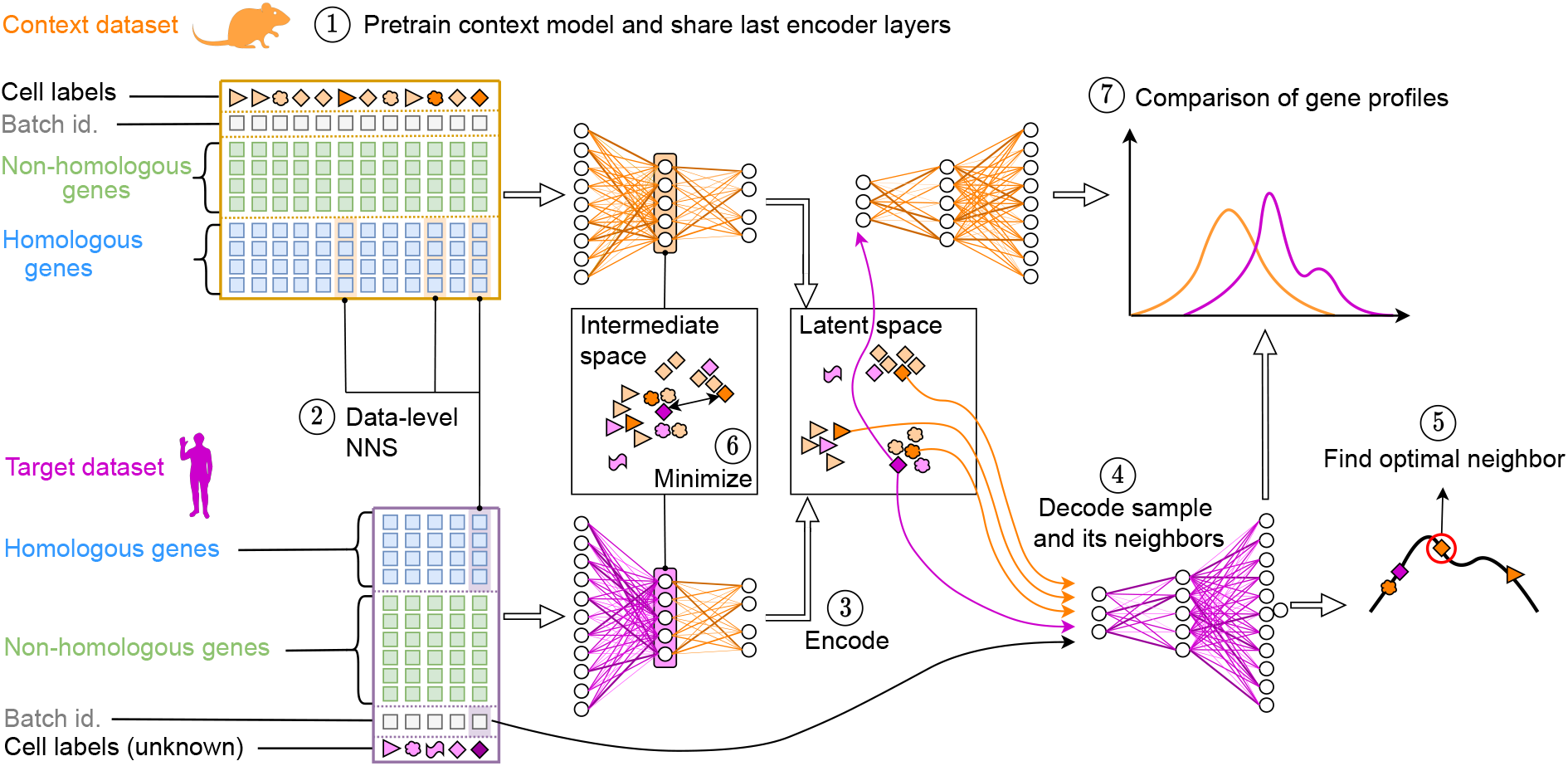
Graphical representation of the scSpecies workflow. Step 1: Encoder and decoder neural networks are trained on the dataset of the context species. Weights of the last encoder layers are incorporated into the encoder model for the target species. Step 2: A nearest neighbor search is performed on shared genes of the context and target dataset. This identifies a set of *k* context neighbors for every target cell. Step 3: Cells of the target dataset are encoded into the latent space. For cells with high agreement among cell labels of their neighbors, we retrieve the latent variables of their neighbors. Step 4: The latent values of their *k* neighbors are passed to the decoder together with the human batch label. Step 5: The optimal candidate among the *k* neighbors is chosen as the cell with highest log-likelihood. Step 6: The distance between the optimal candidate and the intermediate representation of its target cell is minimized. Step 7: After training, normalized gene expression profiles can be compared by decoding latent variables with both decoder networks. Also, labels can be transferred using the aligned latent representation.

In a first step, our proposed approach pre-trains an scVI model on the context dataset. Afterwards, the last encoder layers are transferred into a second scVI model for the target species. The aim of this architecture transfer is to share learned information within the network weights between datasets and species. During subsequent fine-tuning, the shared weights remain frozen while all other weights are optimized. Unlike existing architecture surgery approaches, we align architecture in a reduced intermediate feature space instead of at the data level. This approach is inspired by the notion of mid-level features from computer vision [22, 23]. These represent abstractions of the input image learned by neural networks in their intermediate layers. Mid-level features combine individual elements into more general structures, like contours, specific shapes, or parts of objects. Transfer learning approaches then retrain the last layers to transition these intermediate representations into task-specific network outputs for different datasets. [24] Unlike images, scRNA-seq datasets lack ordered patterns as gene expression vectors can be permuted without changing their information content. Still, the first encoder layers translate dataset-specific features, like influences of experimental batches or interactions between observed genes, into a higher abstraction level. (Figure 5) The resulting representation may correspond to more fundamental cell properties that are less perceptible to noise and systematic differences between species.

Still, the first encoder layers translate dataset-specific features, like influences of experimental batches or interactions between observed genes, into a higher abstraction level. (Figure 5) The resulting representation may correspond to more fundamental cell properties that are less perceptible to noise and systematic differences between species.

To connect the new encoder layers with the pre-trained structure, we identify sets of similar cells through a nearest neighbor search performed on homologous genes. Afterwards, scSpecies minimizes the distance between a target cell’s mid-level representation and a suitable candidate from its set of neighbors. The model determines the most suitable context cell as the candidate whose decoded latent representation yields the highest log-density value at the location of the target cell within the decoder’s distribution. To counter misclassifications, we align mid-level features only for target cells whose context neighbors have high agreement in their cell labels.

During model fitting, we thus encode similarity information both at the original data level and at the level of learned features. The aligned latent space then captures cross-species similarity relations based on the fitted model, which facilitates information transfer across species.

### 3.1 scSpecies aligns architectures across species

We applied the scSpecies workflow to three mouse-human dataset pairs containing liver cells, white adipose tissue cells, and immune response cells to glioblastoma.

We visually examined alignment through UMAP coordinates [25] of the combined latent variables of dataset pairs (Figure 2). The 2D representation showed biological meaningful alignment of cells. Cell types without context counterparts aligned with related cell types or formed distinct clusters.

**Figure 2:**
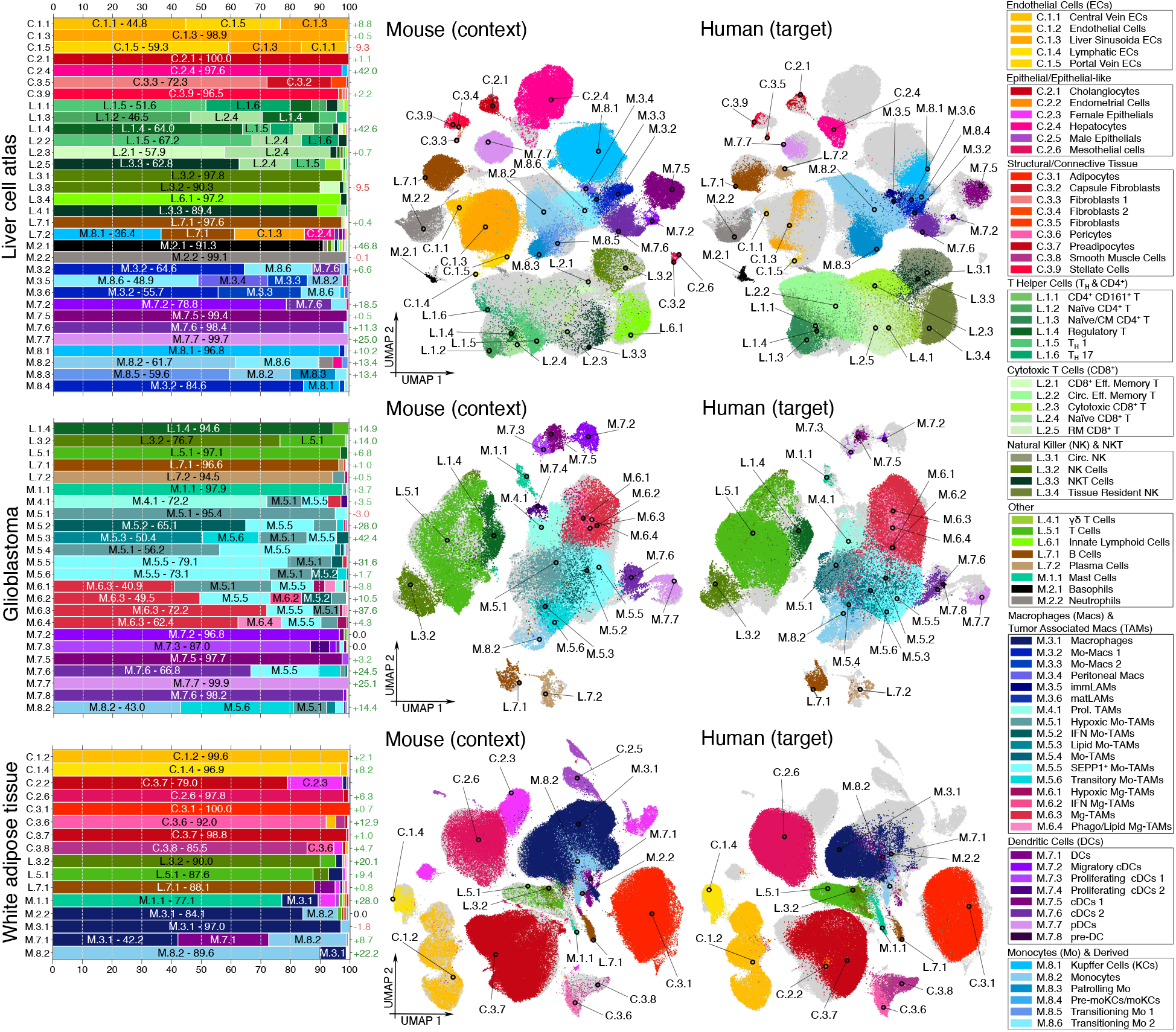
Visualization of the aligned representations for three dataset pairs obtained by training sc-Species with a set of 25 neighbors. We color cells by fine cell type labels for the liver and glioblastoma datasets, and by coarse cell labels for the adipose tissue dataset. On the left, bar plots indicate cell label transfer accuracy through a nearest neighbor search in the aligned latent space. The left y-axis labels indicate cell type codes corresponding to human cell labels. These codes are referenced in the legend. The bars contain the frequency of assigned mouse cell labels. The results are averaged over five random seeds. The left y-axis labels indicate improvement in accuracy for shared cell types over the data-level nearest neighbor search. Next to the bar plots, UMAP coordinates of the aligned latent representations are visualized. Lymphoid cell types are colored in green and brown, myleoid cell types in blue and purple, and CD45^−^ cell types in red, pink and yellow. Cells from the other dataset are indicated in a light gray.

To facilitate label and information transfer for target cells, we conducted a second nearest neighbor search on the shared latent representation of both datasets. Afterwards, we inferred target cell labels from their set of latent context neighbors by majority vote. For labels at the sub-cell type resolution, accuracy was 73% for liver, 49% for adipose tissue, and 69% for glioblastoma datasets. Misclassifications mostly occurred within biologically related cells belonging to the same overarching cell type. For broader cell type labels, accuracy increased to 92% for the liver, 82% for the adipose tissue, and 80% for the glioblastoma dataset. These values represent significant improvements upon the data-level nearest neighbor search and existing architecture surgery approaches (Table 2). We also calculated the adjusted Rand index and adjusted mutual information and observed improvements in these metrics.

We observed higher increase in label transfer accuracy for cell types with noisy data-level nearest neighbor search searches but clear separation in their pre-trained latent space. For example, the initial neighbor search matched less than half of all human liver basophils (cluster M.2.1) with mouse counterparts. This value improved to over 90% through our method. However, in the adipose tissue datasets, neither the context scVI model nor the nearest neighbor search separated dendritic cells, monocytes, and macrophages. Thus, scSpecies could not separate these cell types either.

Results were consistent over architecture variations and averaged over five random seeds, yet for cell types with noisy neighbor search results, like hepatocytes or portal vein endothelial cells, misclassifications of the whole cell type occurred in one random seed.

We also tested scSpecies in a scenario where the target dataset was small but equally diverse in cell types and batch effects. Specifically, we randomly sampled 5000 cells from the human liver dataset and trained the model to align with the full mouse context dataset. We repeated sampling and training ten times and obtained as metrics for the coarse-fine cell labels an accuracy score of 88% - 68%, which still indicates reasonable performance.

### 3.2 The nearest neighbor search is an important component of scSpecies

We explored the importance of incorporating the nearest neighbor search into scSpecies. (Table 8.4) Without this component, we observed misaligned latent representations and significantly reduced label transfer accuracy. Initializing the inner encoder layers with random, frozen weights yielded similar results to using the pre-trained structure. This implies that without neighbor component, transferred layers were treated like random nuisances.

Training with one neighbor forced the model to align some cells with mismatched counterparts as the approach could not choose from a set of suitable options. We observed meaningful alignment but with reduced performance.

Training with 25 neighbors improved results noticeably on all datasets. To investigate the preferred candidate choice, we tracked cell prototypes during alignment. We created context and target prototypes cells consisting of empirical median gene expression values within a cell type. For each target prototype, we included all context prototypes within its set of candidates and tracked their log-likelihoods during alignment (Figure 10). At onset, the likelihoods for all prototypes were nearly equal. This resulted in alignment driven by chance favoring cell candidates of the most occurring cell label. For cell types with a noisy neighbors set, corrections during later training stages eventually aligned them with appropriate prototypes. This is seen with hepatocytes, migratory cDCs or basophils with a nearest neighbor search accuracy of 56%, 61%, and 45% respectively. Cell types where the neighbor search yielded predominantly incorrect results did not align correctly, like killer T cells and cytotoxic CD8^+^ cells, which had an initial accuracy of only 11% and 1%.

Finally, alignment with a large neighbor set caused neglect of rare cell types, resulting in lower corresponding accuracy scores. Metrics like the adjusted Rand index and adjusted mutual information were comparable or improved, as they do not reflect different cell type labels sizes.

### 3.3 scSpecies can help to better separate latent cell clusters

To investigate the intermediate representations, we compared clustering quality of intermediate representations in unaligned and aligned scVI architectures. We found that clustering based on experimental batches became increasingly mixed as data progressed towards the latent space. In unaligned architectures, the Davies-Bouldin index (DBI) increased from 10 to 21.9 in the mouse context, and from 15.8 to 33.5 in the human liver dataset. Conversely, cell type clusters showed increasingly better separation, resulting in a DBI reduction from 4.6 to 1.6 and from 4.9 to 2.4 for the mouse and human datasets. (Figures 5,6,7).

This phenomenon is caused by the design of scVI, which removes batch influences to enforce a normal distribution in the latent space. Batch patterns are added by the decoder through their provided labels. However, scVI must separate cell types to reconstruct cell characteristics from the latent representation. Yet, certain cell types in the human liver dataset, like hepatocytes, stellate cells, and fibroblasts, are predominantly associated with a single batch label. Consequently, the model inferred cell type information from batch labels, removing biological characteristics from their latent variables. However, these cell types were still separated in intermediate spaces which are not regularized to follow a normal distribution.

Alignment adjusted the target encoder architecture to the well-separated latent mouse context representation. This improved latent cell cluster separation, as measured in a decrease in DBI from 2.4 to 1.8. For white adipose tissue and glioblastoma dataset pairs, clustering improvement was marginal, with a decrease in DBI from 1.7 to 1.6 and from 2.2 to 2 respectively.

We also studied the effectiveness of directly aligning latent representations. Direct latent alignment does not require access to the context model weights. However, we observed a decline in performance metrics across all datasets. This underlines the potential of better alignment within the more information-rich mid-level feature spaces.

### 3.4 scSpecies can align datasets of multiple species

We employed scSpecies to simultaneously align liver cells from mice with fatty liver disease, humans, pigs, monkeys, chickens, and hamsters, using a context dataset of healthy mice. (Figure 3)

**Figure 3:**
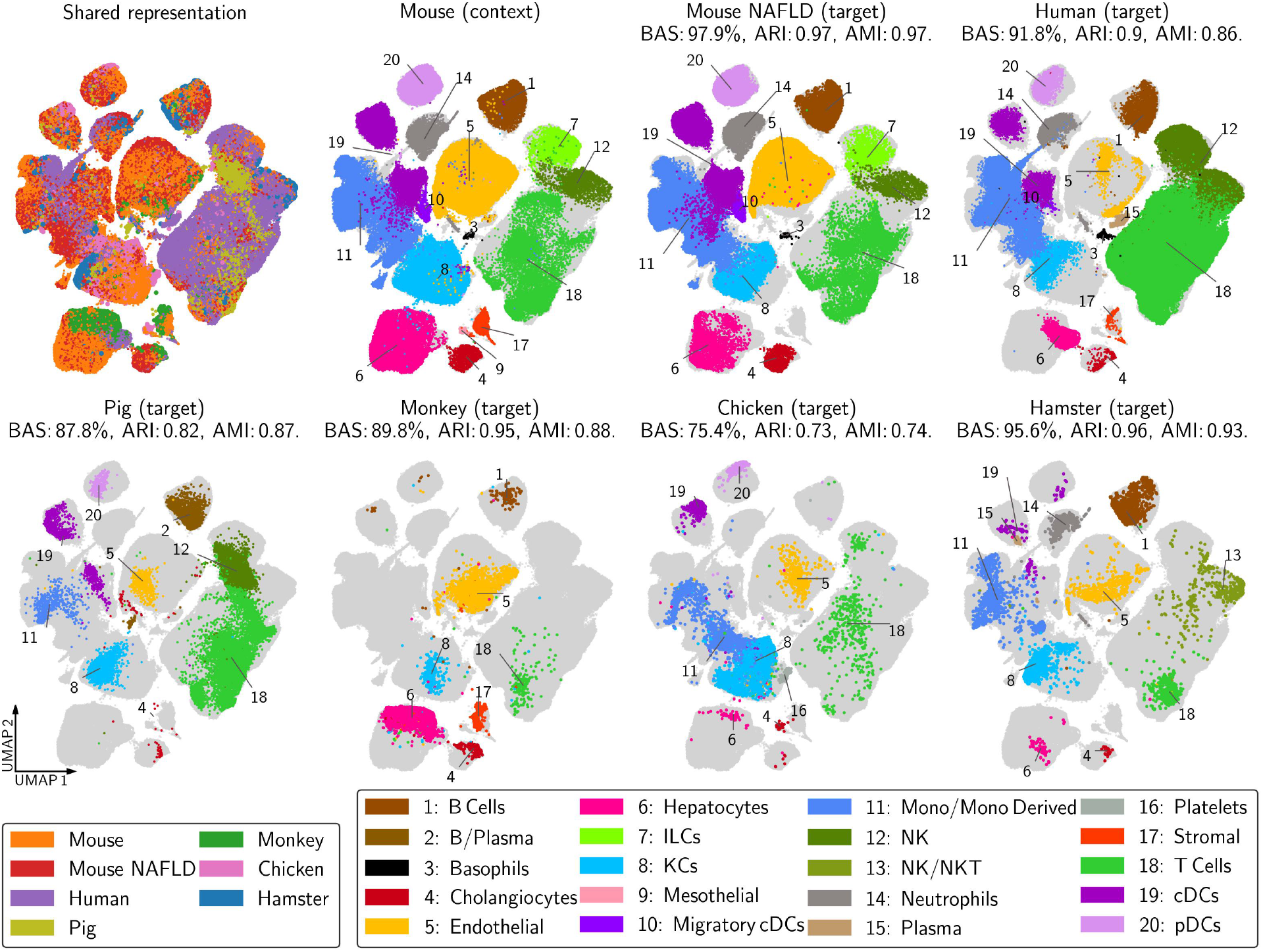
We utilized scSpecies to obtain an aligned liver cell landscape that spans multiple species. The mouse dataset serves as a context for each species.

We successfully obtained aligned latent representations across species, despite less than half of genes having mice orthologs in some datasets.

An intriguing application of scSpecies is the potential to align datasets with very limited gene coverage, or even when there is no overlap in the observed gene set. This can be achieved by aligning both datasets to a comprehensive context dataset with which they both share a common gene set.

However, a limitation of this approach is its inability to align cell types not present in the context dataset. For example, plasma cells, absent in the mouse dataset, were not aligned across the human, pig, and hamster datasets.

### 3.5 scSpecies offers insights into genetic manifestations of cells across species

To better understand the similarities and differences between context and target datasets, e.g. for clarifying in what aspects an animal might be a good model of human biological processes, we extended our analysis from the latent space to the data level. Here, we compared reconstructed gene expression profiles and assigned relevance scores to input genes.

We decoded latent representations using both decoder models to obtain normalized gene expression vectors for each species. These vectors allow us to compare and analyze gene expression profiles of cells that inherit similar underlying biological properties. This analysis benefits from the correspondence between latent representations of both species, which is hard to establish at the data level.

For our investigation, we focused on cell types occurring both in the mouse and human liver datasets. We assessed Log2Fold changes (LFC) in normalized gene expression vectors, which indicates differences of gene expression levels between species. We also calculated the probability of observing genes as differentially expressed when sampling from the latent distribution of a cell type. (Figure 8) Averaging across cell types showed 56% of genes exhibited a LFC ratio above one. Among these, 15% of mouse genes were upregulated and 21% were downregulated compared to their human counterparts in over 90% of decoded cells. With a LFC threshold of two, 24% of genes had a LFC outside this boundary. With a LFC ratio of 0.4, a substantial 82% of genes showed a LFC outside this boundary. These results agree in magnitude with [26], who found an LFC ratio of greater than 0.4 in 78% of genes comparing humans with non-alcoholic liver disease and mice on a high-fat diet.

For white adipose tissue datasets, 50%, and for glioblastoma datasets, 47% of genes exhibited a LFC ratio greater than one.

We compared this against training on context-target datasets pairs of healthy mice and mice with liver disease. Here, only 22% of genes had a LFC change above one. Of those differentially expressed genes, 4% and 5% were upregulated and downregulated in more than 90% of samples. Only 6% of genes had a LFC over two, while 55% of cells showed LFC values above 0.4.

We extended our study by calculating relevance scores using Layer-wise relevance propagation (LRP) [27] (Figure 9). These scores measure each genes contribution to a cells latent value, offering insights into the learned significance of specific genes across different cell types and species. LRP was recently used to explain explain neural network predictions on scRNA-seq data [28].

First, we found no significant difference in relevance scores between non-homologous and shared genes, suggesting that training networks on a reduced gene set omits informative parts of the data.

Second, we found relevance scores to correlate with gene expression levels. For the mice and human liver datasets, we found a Spearman’s *ρ* between the expression level of a gene and their relevance scores of 0.67 and 0.69 and a Pearson correlation coefficient of 0.63 and 0.71. This suggests that differences in gene expression translate into relevant features for the neural networks. A gene with high relevance scores across most cell types was *MALAT1*, which is highly conserved across mammals [29].

## 4 Discussion

We introduced scSpecies, a novel deep learning approach designed to align neural network architectures across species. scSpecies facilitates this alignment even with significant differences between gene sets and expression profiles. A key feature is the integration of a nearest neighbor search within the model, which dynamically aligns biologically close cells during training by leveraging model-based similarity information.

We applied scSpecies to mouse-human dataset pairs of liver, white adipose tissue, and glioblastoma cells, showing successful alignment by verifying biologically close cells occupied similar latent space regions. We demonstrated the viability of the proposed approach even on datasets with a low amount of cell samples.

We exploited the aligned latent representations for label and information transfer and to compare gene expression profiles across species. Therefore, scSpecies not only allows cells to provide context in a unified latent space, but also sheds light on their differential manifestation at the gene expression level. As a limitation, cell types unique to the target dataset tend to get aligned with biologically close cell types in the context dataset instead of being identified as new cells by the approach. Direct metrics to identify new cell types would benefit our method.

While tested with a scVI base model, scSpecies could be extend to multimodal CITE-seq data using multiVI [30].

By aligning datasets from multiple species with a single context dataset, scSpecies offers a shared latent landscape for scRNA-seq datasets across species. Therefore, we envision that our approach will help researchers to contextualize biological properties within their datasets across species.

## 5 Methods

In the following, we represent multidimensional vectors using bold italics and scalar values in regular italics. Dataset elements are indicated with superscript indices, and vector positions with subscript indices. The context dataset is indicated by the subscript *C* and the target dataset by the subscript *T*. Superscripts and subscripts are omitted when they are exchangeable. Random variables are expressed in a sans-serif mathematical font, as in X, Z, L. We represent distributions of random variables with uppercase letters, such as P_Z_, and their probability density functions with lowercase letters, like p_Z_(***z***). Conditional distributions are denoted as P_X|***s***_ := P_X|S=***s***_. In the following, we briefly describe the scVI model, which we subsequently use as a core of our proposed approach.

### 5.1 Single cell variational inference

Consider a dataset 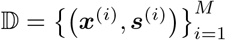 obtained through a single-cell RNA sequencing experiment. The mathematical model behind scVI [21] assumes that gene expression count vectors ***x***, and batch indicator variables ***s***, correspond to observations of random variables X and S. The gene expression data distribution P_X|***s***_ is conditioned on its batch effect S = ***s***. This accounts for technical artifacts during data collection. Within an experimental batch, gene expression vectors are independent and identically distributed samples from P_X|***s***_.

scVI models the data distribution within a parametric family. Building on conditional variational autoencoders [16], a latent variable model is introduced. The random variable Z, corresponding to the representation of a cell in the latent space ℝ^*d*^, is employed to capture biological variability among cells in the dataset. The one-dimensional random variable L with latent space ℝ_*>*0_ accounts for technical variability due to different library sizes. Within the model, data is generated by drawing samples for Z and L from a prior distribution P_Z,L|***s***_. Then, gene expression data is generated by drawing from the sampling distribution P_X|***z***,***l***,***s***_.

The data p.d.f. p_X|***s***_ can be expressed by integrating the joint probability across the latent spaces and then applying the general product rule of probability,

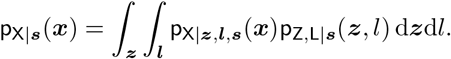

To maximize this integral, scVI performs variational inference on the intractable posterior distribution P_Z,L|***x***,***s***_. Therefore, the posterior probability is approximated by a variational distribution, denoted as Q_Z,L|***x***,***s***_ ≈P_Z,L|***x***,***s***_. Further, scVI applies a mean field approximation, where p.d.fs of both variational and prior distribution are factorized,

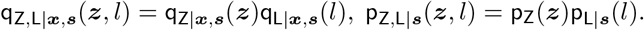

The prior P_Z_ is assumed to be independent of S and fixed as standard normal distribution P_Z_ = 𝒩 (**0, *I***_*d*_). The prior P_L|***s***_ is set as a log-normal distribution 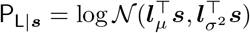. The prior parameters are derived from empirical batch means and variances of the observed log-library sizes. The variational distribution Q_Z|***x***,***s***_ is chosen as a normal distribution 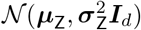, and Q_L|***x***,***s***_ is set as a log-normal distribution 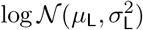.

The parameters for these distributions are determined by two encoder neural networks,

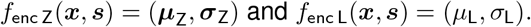

scVI obtains latent variables by sampling from the variational distributions through the reparametrization trick [31].

The sampling distribution P_X|***z***,*l*,***s***_ for generating gene-expression data from a given latent variable is assumed to follow a Gamma-Poisson mixture, resulting in a negative binomial distribution. The corresponding decoder network outputs a denoised gene expression vector that sums to one.

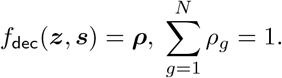

The value *ρ*_*g*_ provides an estimate of the percentage of transcripts in a cell that originate from gene *g*. Gene expression values *x*_*g*_ can be drawn from a negative binomial distribution NB(*lρ*_*g*_, *θ*_*g*,***s***_) parameterized by mean *lρ*_*g*_ and dispersion *θ*_*g*,***s***_. The dispersion parameter is constant for every gene across cells of batch ***s***. To address the potential issue of dropout, a zero-inflated negative binomial distribution can be used to model count data. The dropout probability parameter ***π*** is also obtained from the decoder network. The weights of the three neural networks and the parameters *θ*_*g*,***s***_ are optimized simultaneously by empirically estimating and maximizing the ELBO function

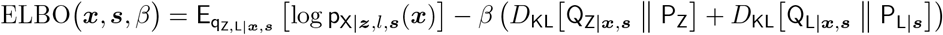

on mini batches 𝕄 ⊂ 𝔻.

## 6 The scSpecies approach

We consider a scenario involving two scRNA-seq datasets,

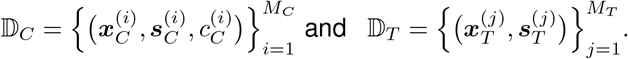

Their data points consist of gene expression measurements ***x*** and batch indicator variables ***s*** from a context species *C* and a target species *T*. Furthermore, context count vectors are clustered into distinct groups based on cell type labels *c*_*C*_, whereas target labels *c*_*T*_ are unknown.

The count vectors from both datasets share a gene subset ***h*** comprising count values from homologous genes,

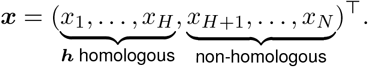

The number of non-homologous genes can differ in both datasets, either because a gene has no ortholog in the genome of the other species or because its is not observed within the dataset. Therefore, gene expression vectors can be of different dimension, *N*_*C*_ ≠ *N*_*T*_.

To map both datasets into a unified latent space, we define separate scVI models for each dataset,

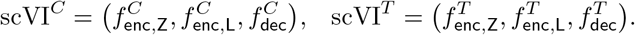

We divide the training procedure for scSpecies into three steps: Training of the context scVI model, followed by an initial data-level nearest neighbor search, and alignment of context and target latent representations.

### 6.1 Pre-training on the context dataset

First, the model scVI^*C*^ is trained on the context dataset by minimizing its negative ELBO function. Following training, the architecture of the encoder network for the latent variable Z is split up into two parts:

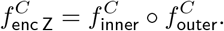

The outer part 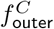 consists of the first *L* layer functions and maps data from the input space 𝒳_*C*_ to an intermediate feature space 𝒯. The inner part, 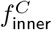, consists of the last *M* layers. It encodes an intermediate representation onto the variational parameters with subsequent reparametrization into the latent space *Ƶ*. We incorporate this inner encoder part into the encoder architecture of scVI^*T*^,

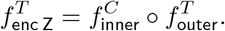

### 6.2 Nearest neighbor search

As the first layers are replaced, the target model scVI^*T*^ cannot pick up on the learned structure within its later encoder layers when starting from a random initialization. Therefore, we incentivize alignment of intermediate target representations with intermediate features of similar context cells. This leads to an aligned latent space as layer weights mapping from the intermediate space to the latent space are not updated. To quantify similarity and establish a direct correspondence between cells of context and target dataset, we perform a nearest neighbor search on the shared homologous gene subset ***h***. The nearest neighbors serve as a set of candidates for every target cell from which the model can choose a best fit to align their intermediate representations during the last training phase.

The nearest neighbor search identifies an index set 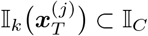 of *k* nearest neighbors for every target gene count vector 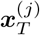. I.e., for every context cell with index 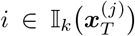 has to hold that a chosen measure of association^1^ between the homologous gene counts, 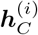 and 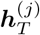, is lower than for elements outside the set:

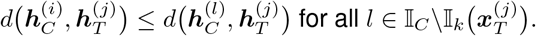

Common metrics or distance functions can be used as a measure of association *d* to compare count values of single-cell data. Some popular choices have been investigated in [32]. We utilize cosine similarity, measuring the cosine of the angle between log1p-transformed count vectors, as it is fast to calculate even on datasets containing numerous samples:

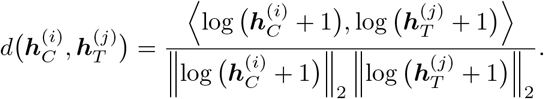

The data-level nearest neighbor search can also be used to assign preliminary labels. We count the multiplicity of cell labels for all context neighbors and assign, as a preliminary label prediction, the most occurring label,

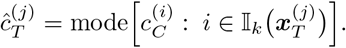

As the data-level nearest neighbor search is noisy, we additionally assign accuracy scores based on the occurrence of a cell label prediction 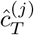.

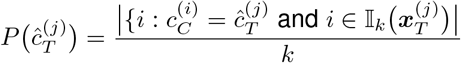

A higher accuracy score indicates lower noise, as there is high agreement among cell labels of the context neighbors. During following alignment, only target cells exhibiting high accuracy scores are considered for alignment in the intermediate space. We collect the indices of all target cells whose accuracy scores 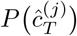 of their predicted cell label *l* are higher than the quantile *Q* at niveau *p*,

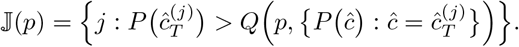

### 6.3 Aligning the intermediate and latent representations

During alignment, the weights of the pre-trained encoder part 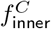 are not updated. To guide the model towards leveraging the learned structure, scSpecies aligns intermediate representations with high accuracy scores

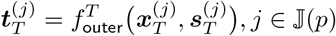

with a representation of a suitable context neighbor representation

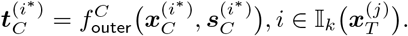

This is facilitated by minimizing the squared Euclidean distance.

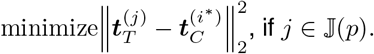

The optimal choice *i*^*^ ∈ 𝕀_*k*_ for minimization among the *k* candidates is dynamically determined during the alignment phase: First, we obtain a set of latent context neighbor variables for the target cells considered during alignment,

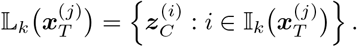

As the context model weights are not updated, the intermediate representations 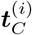 and variational parameters 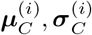, used to obtain the latent variables, can be pre-calculated and stored. These latent variables 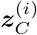 are then decoded with the batch indicator variable 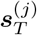 of their target cell. The decoder output and target library size 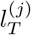 parameterize a sampling distribution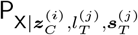, which is used to calculate log density values for every candidate. The candidate whose latent representation results in the highest log density value at 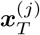 is chosen as optimal neighbor candidate:

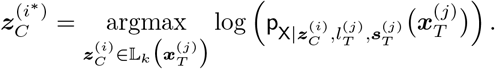

Using this procedure, it is possible to assign a context neighbor with a fitting cell type if at least one candidate with this cell type is found in this set. The training criterion for the model scVI^*T*^ on the target dataset for a data point is

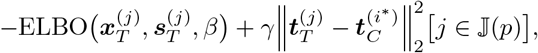

where [*j* ∈ 𝕁(*p*)] is the Iverson Bracket that takes value 1 when an index of a target cell *j* is in 𝕁(*p*), and 0 otherwise. This holds true for cells that exhibited a high degree of agreement during the data-level nearest neighbor search. As minimization in the intermediate space is only incentivized for cells with these indices, the remaining cells within a mini-batch are grouped around them in a way that minimizes the nELBO of the scVI model.

The scalars *γ, β* ≥ 0 weighing different parts of the loss function, the quantile niveau *p* ∈ [0, 1] and number of nearest neighbors *k* ∈ ℕ are hyperparameters.

### 6.4 Transferring cell states and cell types

The aligned latent representations 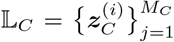 and 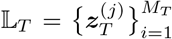 can be analyzed for similarities and differences. For example, their dimensionality can be further reduced into two dimensions using a dimension reduction algorithm like UMAP [25]. To remove the random influence of the latent sampling process, we calculate UMAP coordinates using the variational mean parameters ***µ***.

We can transfer cell labels or cell states from the context to target species by performing a second neighbor search on aligned latent representations. A suitable measure of association is the learned log-density, as it considers the learned manifold of the latent space:

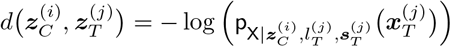

We transfer the most common cell type among the top *k* candidates to the target cell.

### 6.5 Comparison of gene profiles

To perform a comparison of gene expression profiles between cells of context and target dataset, we tailor the methods outlined in [33] and [34] to scSpecies. For a latent variable ***z***, we obtain normalized gene expression profiles by decoding it with both decoder networks and averaging over all possible batches 𝕊:

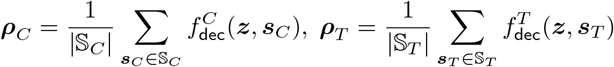

Differences in gene expression profiles can be analyzed for homologous genes, for example, by calculating the log2-fold change (LFC)

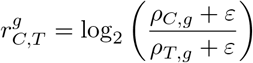

For genes *g* with low expression levels in both species but still high differences, the offset *ε* ensures the associated LFC maintains a low order of magnitude. We modify the decoder output layers to avoid artifacts from the softmax function. These artifacts can arise due to highly expressed non-homologous genes or due to different data dimensions. We apply the softmax function to homologous and non-homologous genes separately to obtain

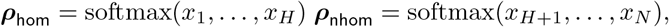

where *N* is the dimensionality of the gene expression vector and *H* the number of homologous genes. Afterwards, both vectors are scaled so that they sum to one,

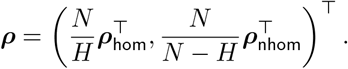

Following [34], for a cell type C = *c*_*C*_ we calculate a mixture distribution of latent states.

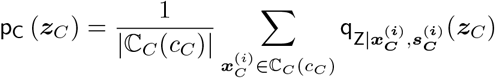

The set ℂ_*C*_(*c*_*C*_) is the set of cells with label *c*_*C*_ with removed outliers. These outliers are identified by estimating the covariance matrix from variational mean samples ***µ***_*C*_. Cells whose variational mean falls outside the 90%-confidence ellipse described by the covariance estimate are removed. A LFC distribution of homologous genes for cell types present in both datasets can be estimated by sampling latent variables from P_C_ and computing the corresponding LFC values 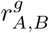. We calculate the median of the empirical LFC distribution as well as the probability 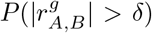 of observing a LFC in gene *g* higher than level *δ >* 0.

### 6.6 Layer-wise relevance propagation

In the following, we briefly describe Layer-wise Relevance Propagation (LRP) [27]. LRP explains the output *f* (***x***) of a neural network *f* by decomposing it into local contributions of input nodes *x*_*i*_, called relevance scores *R*_*i*_(*x*_*i*_). [27]. These relevance scores serve as a measure of each input’s influence on the network’s output: positive scores (*R*_*i*_ *>* 0) signify a positive influence, whereas negative scores (*R*_*i*_ *<* 0) indicate a negative effect. LRP structurally decomposes the function learned by neural networks into a set of smaller, simpler sub-functions of adjacent layers, while ensuring the conservation of relevance scores across the network. This applies locally, where the sum of the relevance score *R*_*i*_ is conserved across two successive layers of the neural network, and globally between the resulting relevance score for each input node *x*_*i*_ and the output *f* (**x**) of the model [27].

Considering a neural network with ReLU activation function, the output *a*_*k*_ of a neuron is given by the input *a*_*j*_ of the previous layer and their connected weights *w*_*jk*_ of the neurons by

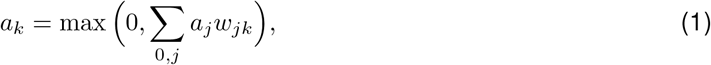

including the bias with *a*_0,*k*_ = 1. The relevance scores *R*_*k*_ describe the contribution of each neuron activation *a*_*j*_ to *a*_*k*_. They can be computed by the LRP-*γ* rule through

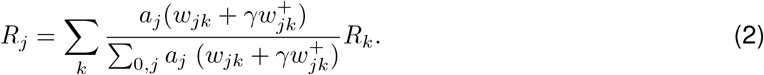

Here, 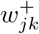 are the positive weights, while *γ* controls how much these positive contributions are emphasized [35]. LRP methodology aligns with the principles of Deep Taylor Decomposition, which breaks down and redistributes the network’s output function *f* (***x***) layer by layer through Taylor series expansions. This decomposition allows for the derivation of various LRP rules tailored to the network architecture and the specific function being analyzed [36]. To compute relevance scores for context and target gene expression vectors ***x***_*C*_, ***x***_*T*_ we propagated the relevance of their latent variational mean parameters ***µ***_*C*_, ***µ***_*T*_ through the corresponding encoder network. We aggregate relevance scores through averaging over latent dimensions and data points of a cell type. A direct comparison of scores between species is complicated by the influence of non-homologous genes and batch-effects on the relevance scores of homologous genes through the conversation property. Rather, ranked lists of genes by scores can be compared across species.

## 7 Metrics

### 7.1 Balanced accuracy score (BAS)

We measured label transfer accuracy by determining the percentage of cells correctly labeled through the nearest neighbor searches on the data level and in the aligned latent space. The calculation was averaged over all cell type classes present in both context and target datasets:

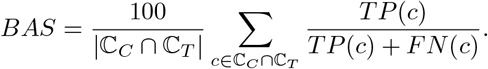

*TP*(*c*) denotes the absolute number of correctly labeled cells within cell label *c* and *FP*(*c*) the number of misclassifications. Cell types unique to the target dataset are not considered by this metric.

### 7.2 Adjusted Rand index (ARI)

We utilized the Rand index (RI) to measure similarity between the clustering given by the true target cell labels 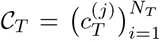 and assigned cell labels 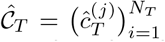 through the nearest neighbor search.

The RI quantifies the agreement of cell pairs being assigned to the same or different cell types in both clusterings [37].

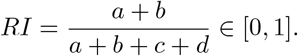

In this formula,

*a* = Number of cell pairs that have the same label in 𝒞_*T*_ and 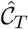,

*b* = Number of cell pairs that have different labels in 𝒞_*T*_ and 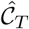,

*c* = Number of cell pairs with the same label in 𝒞_*T*_ but a different one in 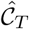,

*d* = Number of cell pairs with a different label in 𝒞_*T*_ but the same one in 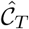,

The ARI enhances the Rand index by adjusting for chance, considering the expected similarity of all pairings.

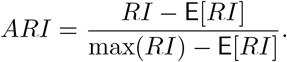

An ARI of 1 indicates perfect agreement between true and predicted cell labels, and values near 0 indicate random label assignments. The ARI considers cell types unique to both datasets and unique ones to the context and target dataset.

### 7.3 Adjusted mutual information (AMI)

We used the mutual information score (MI) to quantify the shared information about the true labels provided by the predicted labels [37]:

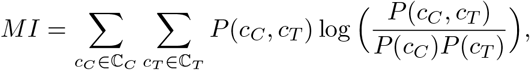

where *P* (*c*_*C*_) and *P* (*c*_*T*_) are the probabilities that a cell is assigned to label *c*_*C*_ in clustering 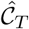 and *c*_*T*_ in clustering 𝒞_*T*_, respectively. *P* (*c*_*T*_, *c*_*C*_) is the probability that a cell with true label *c*_*T*_ is assigned to cell type *c*_*C*_. Similar to the ARI, the adjusted mutual information score corrects for chance, ensuring that the score genuinely reflects agreement beyond random expectations. A score of 1 indicates perfect correlation, meaning an exact match between predicted and true cell labels. A score near 0 implies that the quality is equivalent to that of random clustering. The formula for the AMI is given by

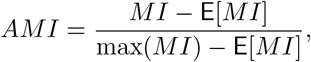

### 7.4 Davies-Bouldin index (DBI)

We evaluated clustering of intermediate representations by cell types and experimental batch effects using the Davies-Bouldin index [38]. Let 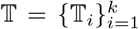 be the intermediate representation of context or target dataset at an arbitrary encoder layer. The representation is clustered either by true cell clusters or experimental batch information. For each cluster, let avg(𝕋_*i*_) be the average distance of all elements in 𝕋_*I*_ to the centroid of 𝕋_*i*_, representing the within-cluster dispersion. Let *d*(𝕋_*i*_, 𝕋_*j*_) be the Euclidean distance between centroids of clusters 𝕋_*i*_ and 𝕋_*i*_, representing the between-cluster separation. The Davies-Bouldin index is then defined as:

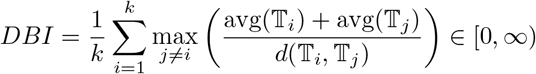

The DBI assesses the average similarity between clusters, defined by the ratio of within-cluster dispersion to between-cluster separation. The index is bounded below by 0, and lower scores indicate better clustering, as they imply compact, well-separated clusters.

## 8 Code availability

Our model is implemented in PyTorch 2.1. The preprocessing scripts to obtain the datasets and the code to reproduce our results can be accessed at https://github.com/cschaech/scSpecies. We recommend to use a device equipped with an NVIDIA GPU.

### 8.1 Hyperparameters

All models were trained with the same network architecture. Gene expression was modeled using a zero-inflated negative binomial distribution with constant dispersion for genes within an experimental batch. We chose a 10-dimensional latent space and a 300-dimensional intermediate space and mapped to and from these spaces with network architectures listed in Table 1. We trained models for 30 epochs on datasets with more than 10,000 cells and 60 epochs on datasets with less observed samples. Network parameters were updated with the ADAM optimizer [39] using standard hyperparameters and a batch size of *M* = 128.

**Table 1:** The network architecture used for all models. *N* denotes the gene expression data dimension, and *S* the number of batch effects. Layer functions contain an affine linear transformation, followed by layer normalization (LN), ReLU activation functions which are clipped to the interval [0, 6], and dropout layers with a dropout rate of *p* = 0.1. Latent representations are obtained from the variational mean and scale encoder model output via the reparametrization trick.

| Model | Layer | In | Architecture | Out |
| --- | --- | --- | --- | --- |
| $f_{\text{outer}}$ | 1 | $N + S$ | Linear, LN, ReLU, Dropout | 300 |
| $f_{\text{inner}}$ | 1 | 300 | Linear, LN, ReLU, Dropout | 200 |
| | 2 | 200 | Linear $\rightarrow 2 \cdot 10$ Rep. trick | 10 |
| $f_{\text{enc L}}$ | 1 | $N + S$ | Linear, LN, ReLU, Dropout | 200 |
| | 2 | 200 | Linear $\rightarrow 2 \cdot 1$ Rep. trick | 1 |
| $f_{\text{dec}}$ | 1 | $10 + S$ | Linear, LN, ReLU, Dropout | 200 |
|  | 2 | 200 | Linear, LN, ReLU, Dropout | 300 |
| | 3 | 300 | Linear, (Softmax, Sigmoid) | $2N$ |
| | $\theta_{g,s}$ | $S$ | Matrix multiplication | $N$ |

**Table 2:** The datasets employed for evaluating scSpecies use mice as context species *C*. The number *H* of homologous genes of context and target dataset are listed in the third column. Furthermore, all datasets are annotated with cell type labels, both at coarse and fine levels. The amount of distinct labels are detailed in the ‘Number of cell labels’ columns. Additionally, the amount of shared cell labels with the context dataset, are indicated in parentheses.

| Dataset | Organism | Shared genes<br>$H$ | Cells<br>$M$ | Batches<br>$S$ | Number of cell types | |
| --- | --- | --- | --- | --- | --- | --- |
|  |  |  |  |  | Coarse | Fine |
| Liver | *C* Mouse | 4 000 | 165 680 | 34 | 15 (15) | 36 (36) |
|  | *T* Mouse NAFLD | 2 860 | 91 787 | 22 | 14 (14) | 28 (22) |
|  | *T* Human | 1 808 | 146 839 | 30 | 15 (14) | 32 (20) |
|  | *T* Human small | 1 808 | 5 000 | 30 | 15 (14) | 32 (20) |
|  | *T* Pig | 1 694 | 21 907 | 2 | 9 (8) | unknown |
|  | *T* Monkey | 1 293 | 8 483 | 2 | 7 (7) | unknown |
|  | *T* Chicken | 1 197 | 7 456 | 2 | 9 (7) | unknown |
|  | *T* Hamster | 1 662 | 5 955 | 2 | 11 (9) | unknown |
| White fat | *C* Mouse | 4 000 | 192 470 | 26 | 17 (17) | 47 (47) |
|  | *T* Human | 1 937 | 137 306 | 24 | 16 (15) | 44 (37) |
| Glioblastoma | *C* Mouse | 4 000 | 46 321 | 6 | 14 (14) | 23 (23) |
|  | *T* Human | 1 823 | 58 560 | 12 | 14 (14) | 24 (22) |

We chose to weigh the KL-Divergence terms with *β* = 0.1 at epoch 1, incrementally increasing their influence to *β* = 1 over 10 epochs. Similarly, the alignment term started with a weight of *η* = 10, which was raised to *η* = 25. The number of nearest neighbors was set to *k* = 25 and the quantile cut-off for alignment was set to *p* = 0.8 across datasets exceeding 10,000 samples. For smaller datasets, we lowered the threshold to *p* = 0.6 to avoid discrimination against scarce cell types. In the latent nearest neighbor search, we pre-computed for each target cell a set of 200 nearest neighbors using the Euclidean distance between the variational mean vectors. Among the 25 cells that resulted in the highest likelihood values, we transferred the most occurring cell label. For differential gene expression analysis, we sampled 10,000 times from the plugin estimator and set the offset variable to *ε* = 10^−6^.

To compute layer-wise relevance scores we retrained the networks with unbounded ReLU activation functions and without layer normalization, as it is difficult for LRP to handle normalization layers. To counteract exploding intermediate values caused by high gene expression values, we trained the model on log1ptransformed values. Omitting layer normalization lead to a slight performance drop of around 2.5% across all performance metrics. We calculated relevance scores using the LRP-*γ* rule with *γ* = 0.15.

We trained both scArches and scPoli on a scVI base model using the scArches package implementation. These models were trained with the same network architecture as scSpecies. We trained both models on homologous genes, as the scArches publication states that zero-filling only produces reliable results when less than 25% of genes are affected [9][See feature overlap between reference and query]. scPoli received training with 10-dimensional batch representations. All other hyperparameters were left at default values.

### 8.2 Datasets

Our model underwent testing on publicly available datasets. (Table 2)

The ‘Liver Cell Atlas’ [17] contains a diverse collection of liver cells from multiple species, including mice (both with and without non-alcoholic fatty liver disease), humans, pigs, monkeys, chickens, and hamsters. We utilized all cells acquired through the scRNA-seq and CITE-seq pipelines. The datasets are available for download at https://www.livercellatlas.org/.

The ‘Single-Cell Atlas of Human and Mouse White Adipose Tissue’ [18] contains gene expression data from human and murine white fat cells. We selected cell samples obtained via single-nucleus sequencing. The data is accessible at https://singlecell.broadinstitute.org/single_cell/study/SCP1376.

The ‘Brain Immune Atlas’ profiles immune response to a grade IV glioma. For humans we selected cells obtained via scRNA-seq of newly diagnosed and recurrent glioblastoma. For mice we selected cells from the immune response to transplanted glioblastoma [19, 20]. The data can accessed via https://www.brainimmuneatlas.org/.

We applied a uniform pre-processing pipeline across all datasets. Initially, the dimension of gene expression vectors was reduced to 4000 most highly variable genes [40]. Then we excluded cells with less than 2% nonzero genes or belonging to extremely scarce batch and cell labels with less than 20 samples. We identified gene homologs using the `HOM_MouseHumanSequence.rpt` gene mapping, available on the MGI Data and Statistical Reports website at http://www.informatics.jax.org/downloads/reports/index.html. To obtain a consistent nomenclature between the datasets some cell labels were renamed. In the liver and glioblastoma datasets, some cells have inconsistent cell type labels. For example, some human liver cells are labeled as neutrophils in the fine and monocytes in the coarse cell label category. We excluded all cells with such a labeling conflict.

## 8.4 Extended Data

**Table 3:**
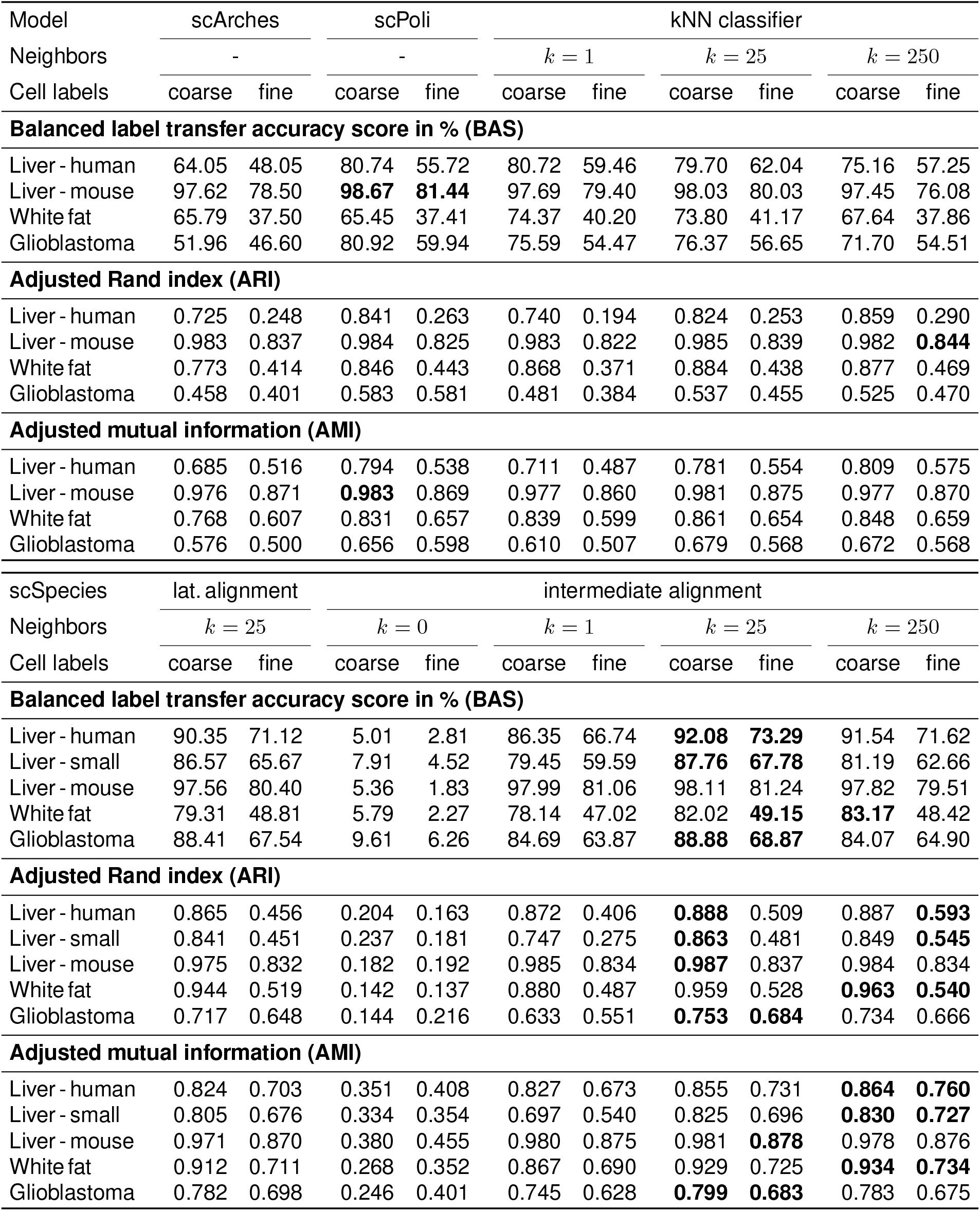
Comparison of model performance on four different datasets. The results are averaged over five random seeds and the best results highlighted by bold font. The results for each dataset are listed for the coarse - fine cell label categories. The upper table contains the results obtained by scArches and scPoli. The kNN columns refer to the results of a data-level *k* nearest neighbor classifier trained on shared homologous genes. The results from scSpecies are listed in the bottom table. The first column corresponds to the results of a scSpecies model where latent representations instead of the intermediate representations are aligned. The column with zero neighbors corresponds to completely omitting the nearest neighbor integration within the model. The column with one neighbor corresponds to omitting learning a suitable neighbor candidate, as the choice is fixed.

| Model | scArches |  | scPoli |  | kNN classifier |  |  |  |  |  |
| --- | --- | --- | --- | --- | --- | --- | --- | --- | --- | --- |
| Neighbors | - | | - | | $k = 1$ | | $k = 25$ | | $k = 250$ | |
| Cell labels | coarse | fine | coarse | fine | coarse | fine | coarse | fine | coarse | fine |
| **Balanced label transfer accuracy score in % (BAS)** |  |  |  |  |  |  |  |  |  |  |
| Liver - human | 64.05 | 48.05 | 80.74 | 55.72 | 80.72 | 59.46 | 79.70 | 62.04 | 75.16 | 57.25 |
| Liver - mouse | 97.62 | 78.50 | **98.67** | **81.44** | 97.69 | 79.40 | 98.03 | 80.03 | 97.45 | 76.08 |
| White fat | 65.79 | 37.50 | 65.45 | 37.41 | 74.37 | 40.20 | 73.80 | 41.17 | 67.64 | 37.86 |
| Glioblastoma | 51.96 | 46.60 | 80.92 | 59.94 | 75.59 | 54.47 | 76.37 | 56.65 | 71.70 | 54.51 |
| **Adjusted Rand index (ARI)** |  |  |  |  |  |  |  |  |  |  |
| Liver - human | 0.725 | 0.248 | 0.841 | 0.263 | 0.740 | 0.194 | 0.824 | 0.253 | 0.859 | 0.290 |
| Liver - mouse | 0.983 | 0.837 | 0.984 | 0.825 | 0.983 | 0.822 | 0.985 | 0.839 | 0.982 | **0.844** |
| White fat | 0.773 | 0.414 | 0.846 | 0.443 | 0.868 | 0.371 | 0.884 | 0.438 | 0.877 | 0.469 |
| Glioblastoma | 0.458 | 0.401 | 0.583 | 0.581 | 0.481 | 0.384 | 0.537 | 0.455 | 0.525 | 0.470 |
| **Adjusted mutual information (AMI)** |  |  |  |  |  |  |  |  |  |  |
| Liver - human | 0.685 | 0.516 | 0.794 | 0.538 | 0.711 | 0.487 | 0.781 | 0.554 | 0.809 | 0.575 |
| Liver - mouse | 0.976 | 0.871 | **0.983** | 0.869 | 0.977 | 0.860 | 0.981 | 0.875 | 0.977 | 0.870 |
| White fat | 0.768 | 0.607 | 0.831 | 0.657 | 0.839 | 0.599 | 0.861 | 0.654 | 0.848 | 0.659 |
| Glioblastoma | 0.576 | 0.500 | 0.656 | 0.598 | 0.610 | 0.507 | 0.679 | 0.568 | 0.672 | 0.568 |

Table 3: Comparison of model performance on four different datasets.
| scSpecies | lat. alignment |  | intermediate alignment |  |  |  |  |  |  |  |
| --- | --- | --- | --- | --- | --- | --- | --- | --- | --- | --- |
| Neighbors | $k = 25$ | | $k = 0$ | | $k = 1$ | | $k = 25$ | | $k = 250$ | |
| Cell labels | coarse | fine | coarse | fine | coarse | fine | coarse | fine | coarse | fine |
| **Balanced label transfer accuracy score in % (BAS)** |  |  |  |  |  |  |  |  |  |  |
| Liver - human | 90.35 | 71.12 | 5.01 | 2.81 | 86.35 | 66.74 | **92.08** | **73.29** | 91.54 | 71.62 |
| Liver - small | 86.57 | 65.67 | 7.91 | 4.52 | 79.45 | 59.59 | **87.76** | **67.78** | 81.19 | 62.66 |
| Liver - mouse | 97.56 | 80.40 | 5.36 | 1.83 | 97.99 | 81.06 | 98.11 | 81.24 | 97.82 | 79.51 |
| White fat | 79.31 | 48.81 | 5.79 | 2.27 | 78.14 | 47.02 | 82.02 | **49.15** | **83.17** | 48.42 |
| Glioblastoma | 88.41 | 67.54 | 9.61 | 6.26 | 84.69 | 63.87 | **88.88** | **68.87** | 84.07 | 64.90 |
| **Adjusted Rand index (ARI)** |  |  |  |  |  |  |  |  |  |  |
| Liver - human | 0.865 | 0.456 | 0.204 | 0.163 | 0.872 | 0.406 | **0.888** | 0.509 | 0.887 | **0.593** |
| Liver - small | 0.841 | 0.451 | 0.237 | 0.181 | 0.747 | 0.275 | **0.863** | 0.481 | 0.849 | **0.545** |
| Liver - mouse | 0.975 | 0.832 | 0.182 | 0.192 | 0.985 | 0.834 | **0.987** | 0.837 | 0.984 | 0.834 |
| White fat | 0.944 | 0.519 | 0.142 | 0.137 | 0.880 | 0.487 | 0.959 | 0.528 | **0.963** | **0.540** |
| Glioblastoma | 0.717 | 0.648 | 0.144 | 0.216 | 0.633 | 0.551 | **0.753** | **0.684** | 0.734 | 0.666 |
| **Adjusted mutual information (AMI)** |  |  |  |  |  |  |  |  |  |  |
| Liver - human | 0.824 | 0.703 | 0.351 | 0.408 | 0.827 | 0.673 | 0.855 | 0.731 | **0.864** | **0.760** |
| Liver - small | 0.805 | 0.676 | 0.334 | 0.354 | 0.697 | 0.540 | 0.825 | 0.696 | **0.830** | **0.727** |
| Liver - mouse | 0.971 | 0.870 | 0.380 | 0.455 | 0.980 | 0.875 | 0.981 | **0.878** | 0.978 | 0.876 |
| White fat | 0.912 | 0.711 | 0.268 | 0.352 | 0.867 | 0.690 | 0.929 | 0.725 | **0.934** | **0.734** |
| Glioblastoma | 0.782 | 0.698 | 0.246 | 0.401 | 0.745 | 0.628 | **0.799** | **0.683** | 0.783 | 0.675 |

**Figure 4:**
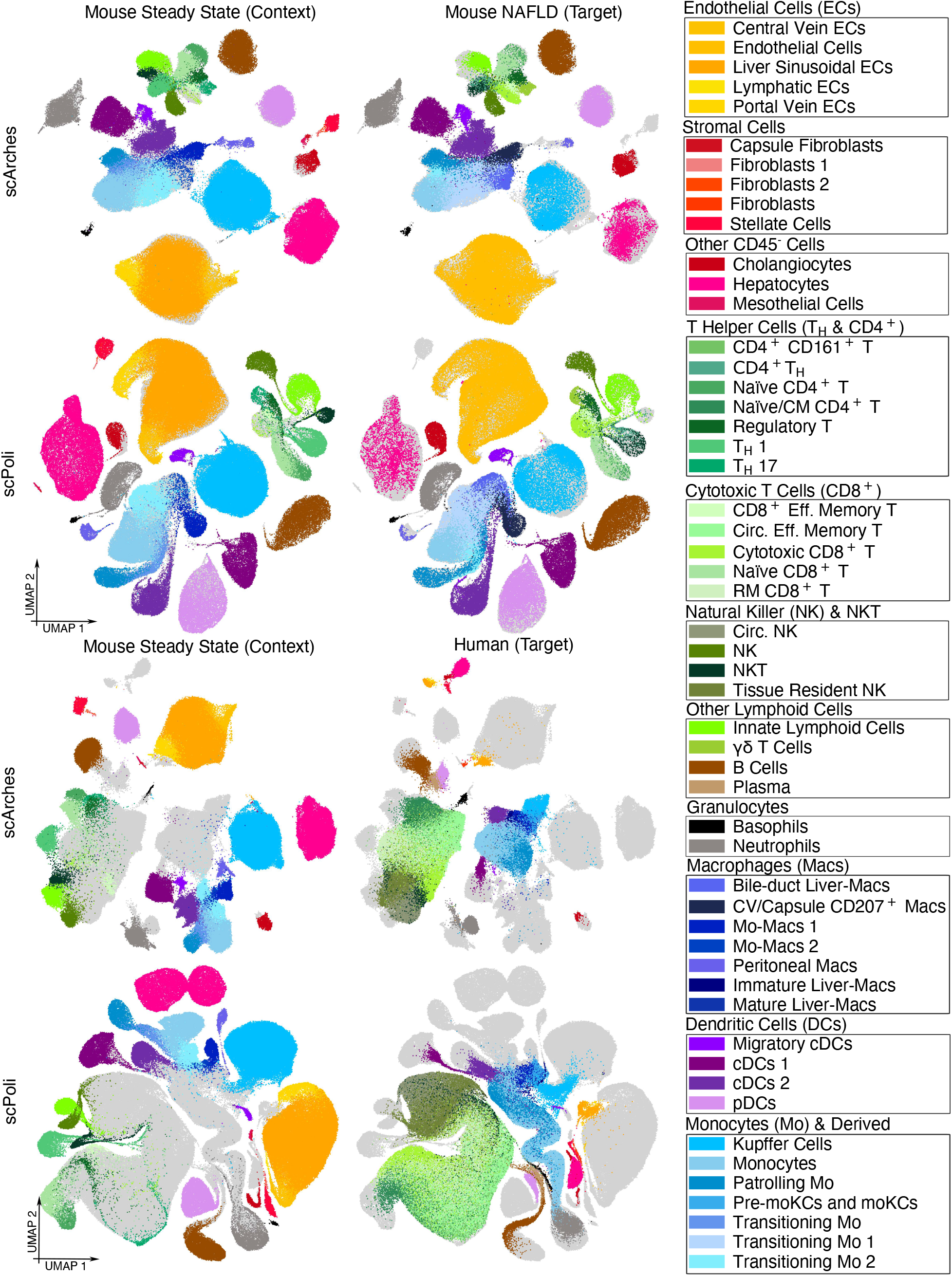
Alignment performance of the architecture surgery-based approaches scArches and scPoli. The four left-hand plots were generated by aligning two mouse liver cell datasets. One dataset contains cell samples from healthy organisms, while the other contains cells from mice with non-alcoholic fatty liver disease. Despite the difference in disease conditions the latent representations are well aligned. The four plots on the right side were obtained by aligning human liver cells with those of healthy mice. Here, both approaches encounter difficulties with cross-species alignment.

**Figure 5:**
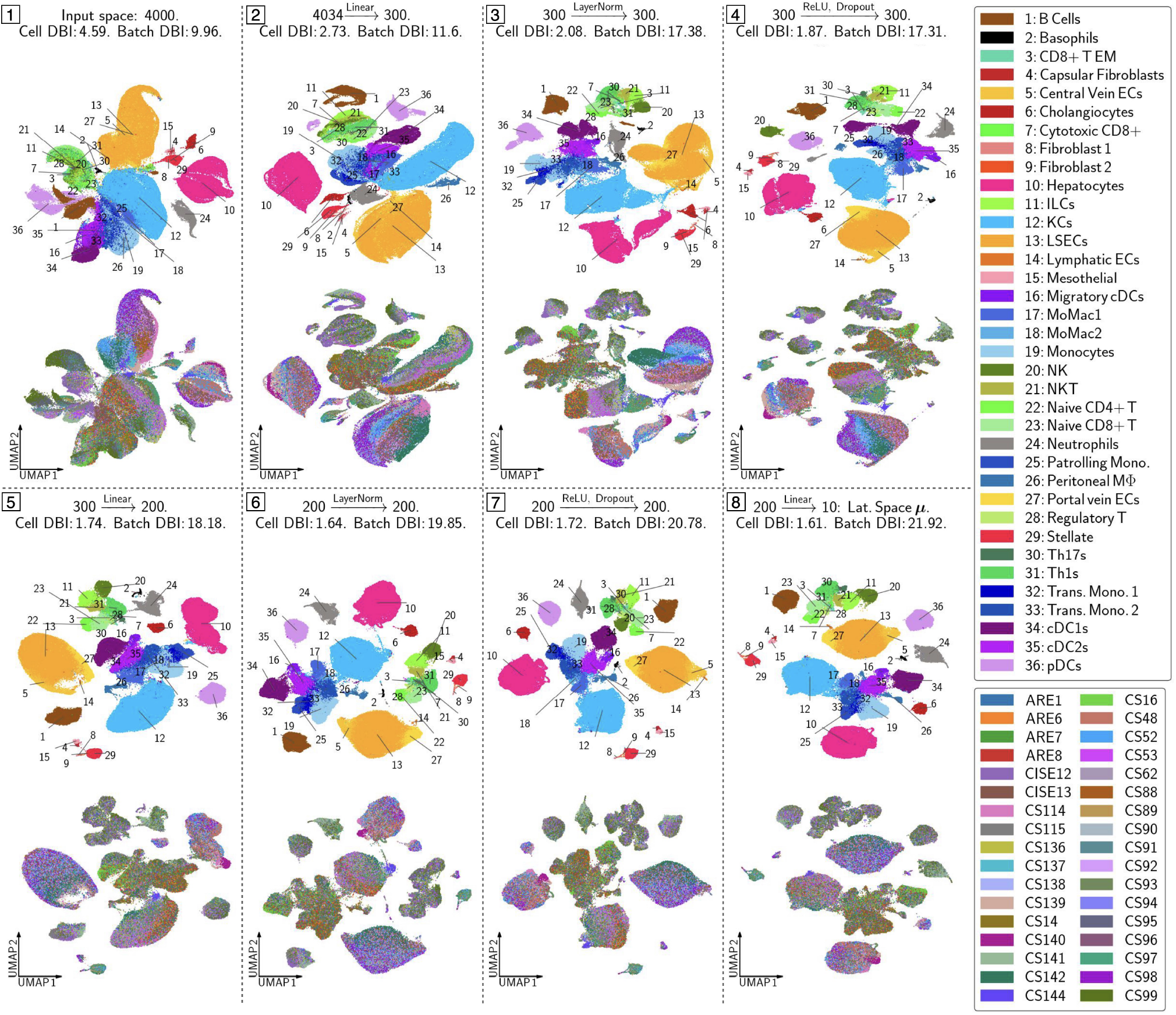
Intermediate spaces of a scVI model applied to the mouse liver context dataset. It details the layer transformations from data space to latent space. Subplot 1 represents the UMAP coordinates of the original dataset, while subplot 8 shows the variational mean vectors in the latent space. Subplots 2–7 depict the UMAP coordinates of the intermediate dataset representation obtained by applying the corresponding layer transformation. Each subplot presents two scatter plots: the upper one showing clusters based on cell labels and the lower one depicting experimental batches. Additionally, the Davies-Bouldin index is used to assess the clustering quality for each subplot.

**Figure 6:**
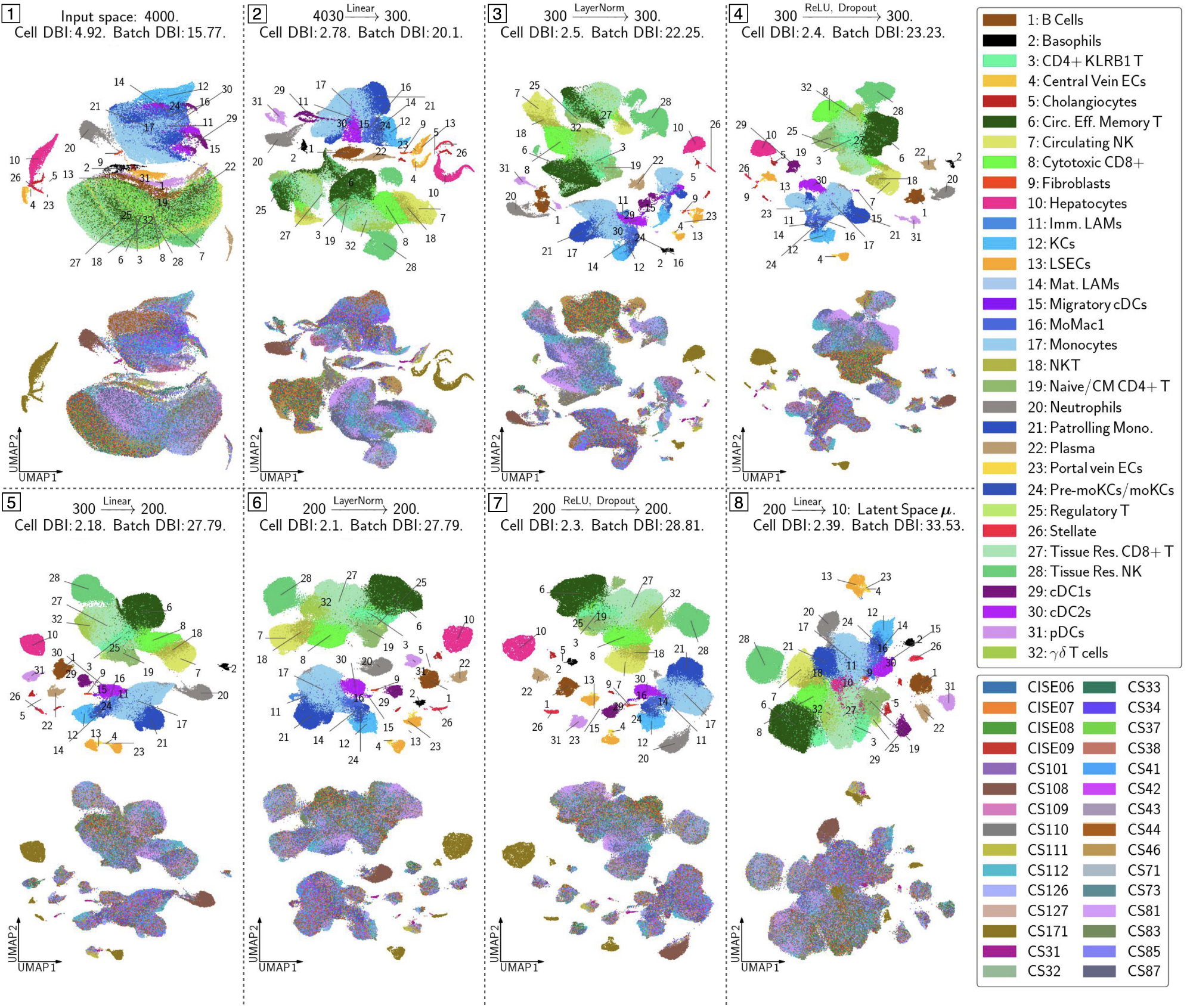
Intermediate spaces of a scVI model applied to the unaligned human liver target dataset. For an explanation of the subplots, see Figure 5.

**Figure 7:**
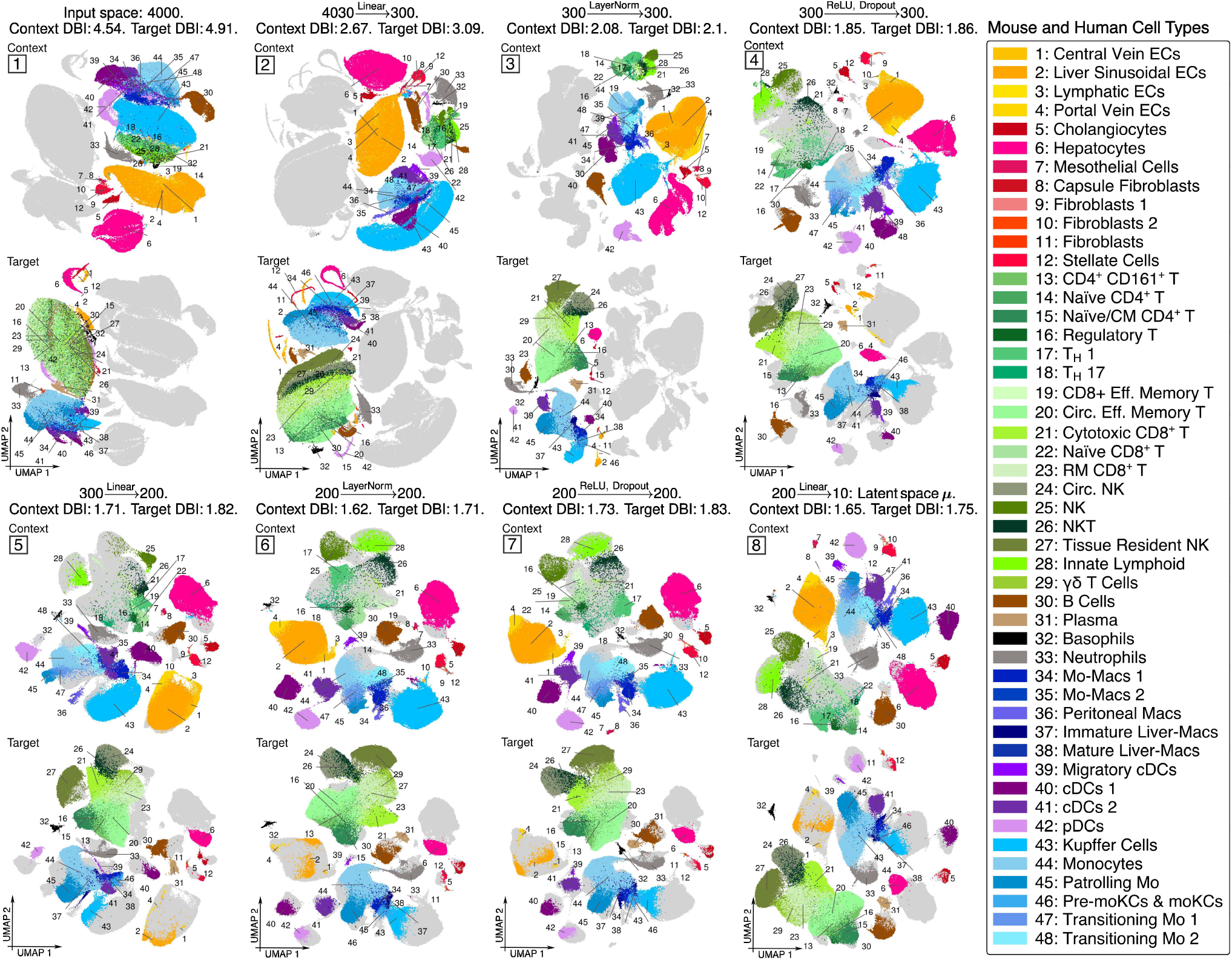
Intermediate spaces of a scSpecies model applied to the mouse-human liver dataset pair. Each subplot presents two scatter plots: the upper one showing context cell label clusters and the lower one depicting the human target cell clusters. Additionally, the Davies-Bouldin index is used to asses clustering quality for each subplot. Alignment of the two datasets is encouraged in subplot 4.

**Figure 8:**
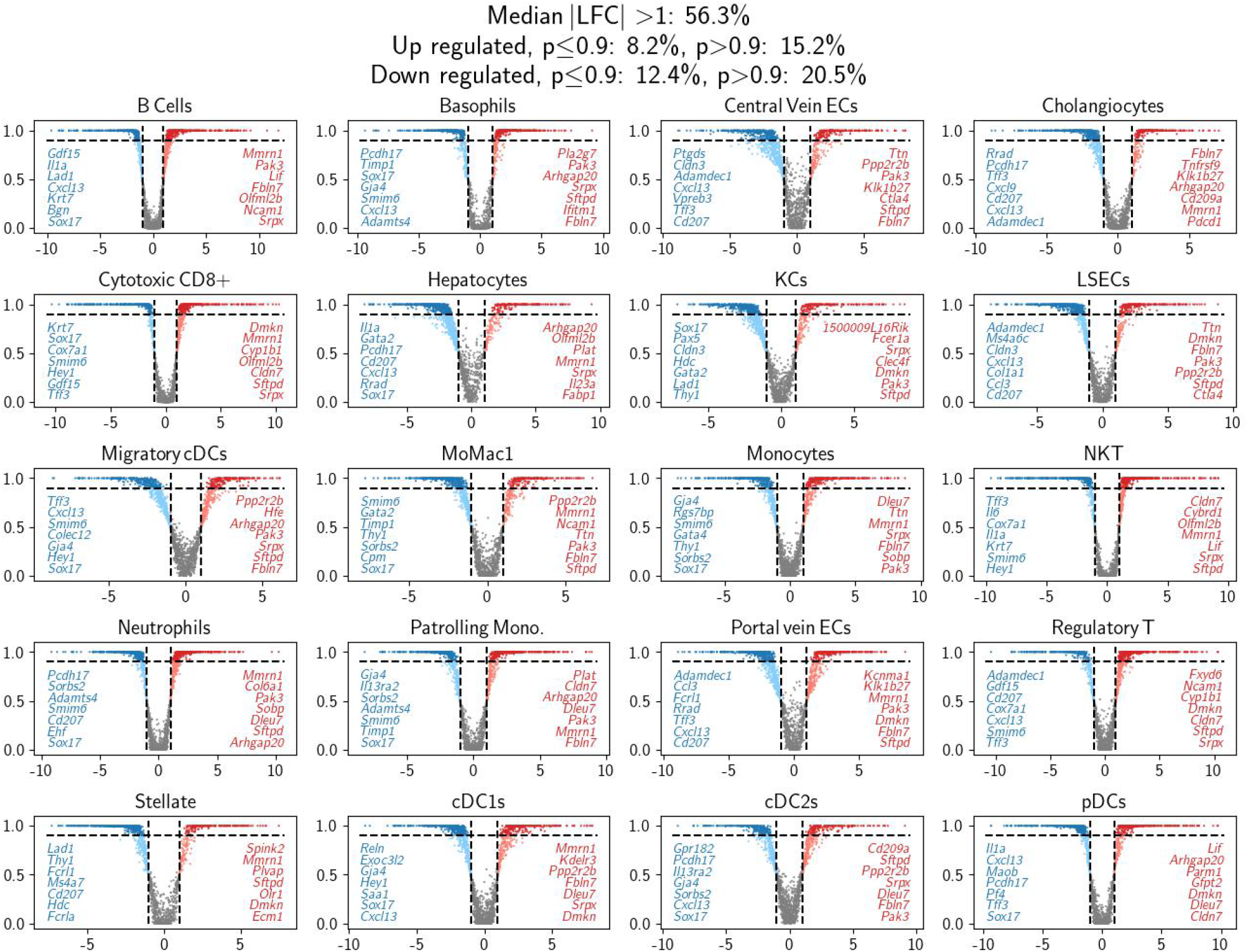
A comparative analysis of gene expression profiles between humans and mice using scSpecies. We computed the median of the empirical log2 fold change distribution, displayed along the x-axis. The y-axis illustrates the likelihood of a gene being differentially expressed in mice versus humans with an LFC exceeding one. The compared cells are decoded from a randomly selected latent value within a latent cell type distribution. The figure highlights the top seven genes in mice that are significantly up-regulated (indicated in red) and the top seven that are notably down-regulated (blue) in comparison to their human equivalents.

**Figure 9:**
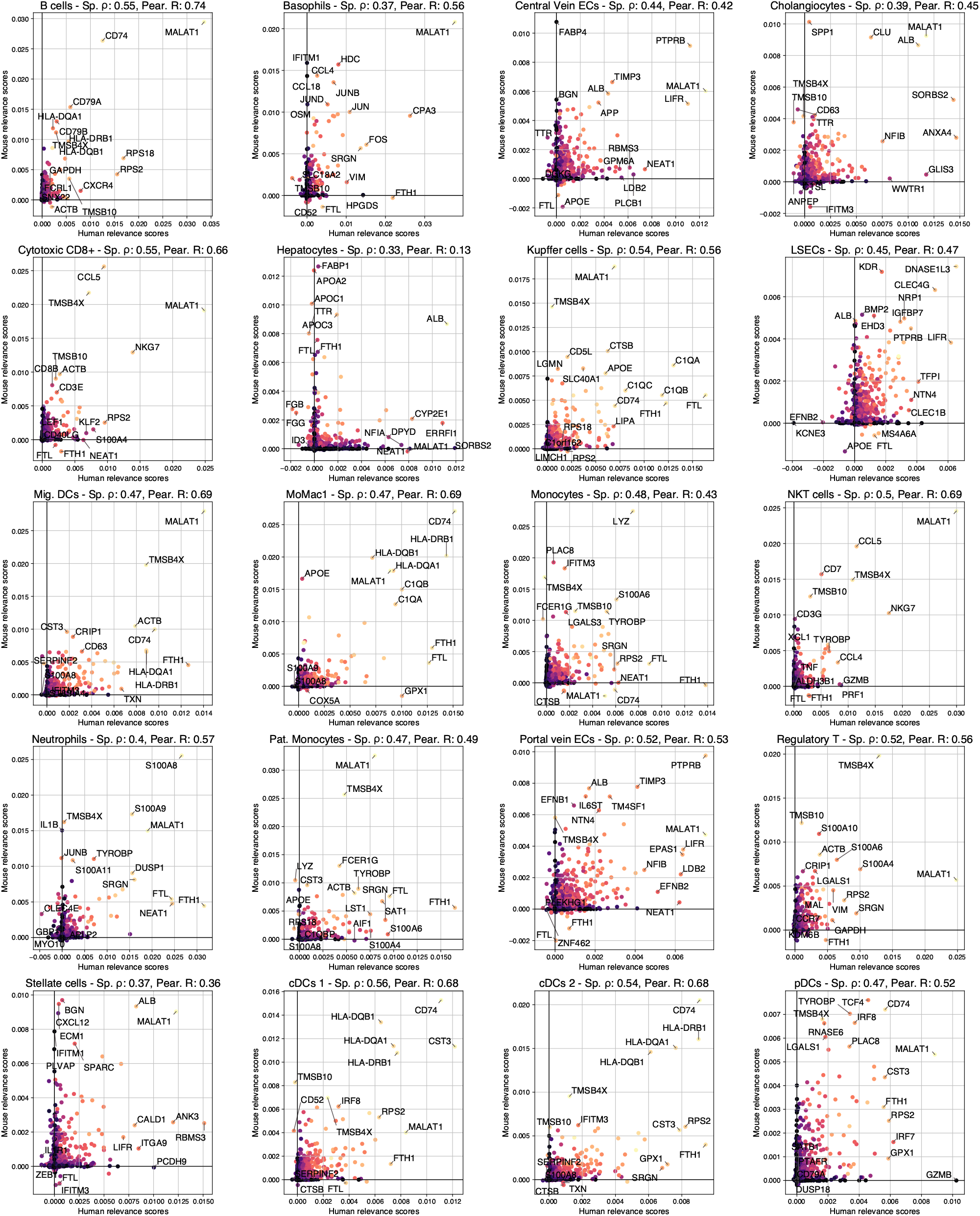
Plots of human and mouse gene LRP scores against each other. Each dot represents a homologous gene. For every cell, Spearman’s *ρ* and Person’s R between human and mice LRP values are given in the axis label. Coloring corresponds to combined products of human and mice gene expression, with values of 0 are colored in dark tones and high values in bright colors.

**Figure 10:**
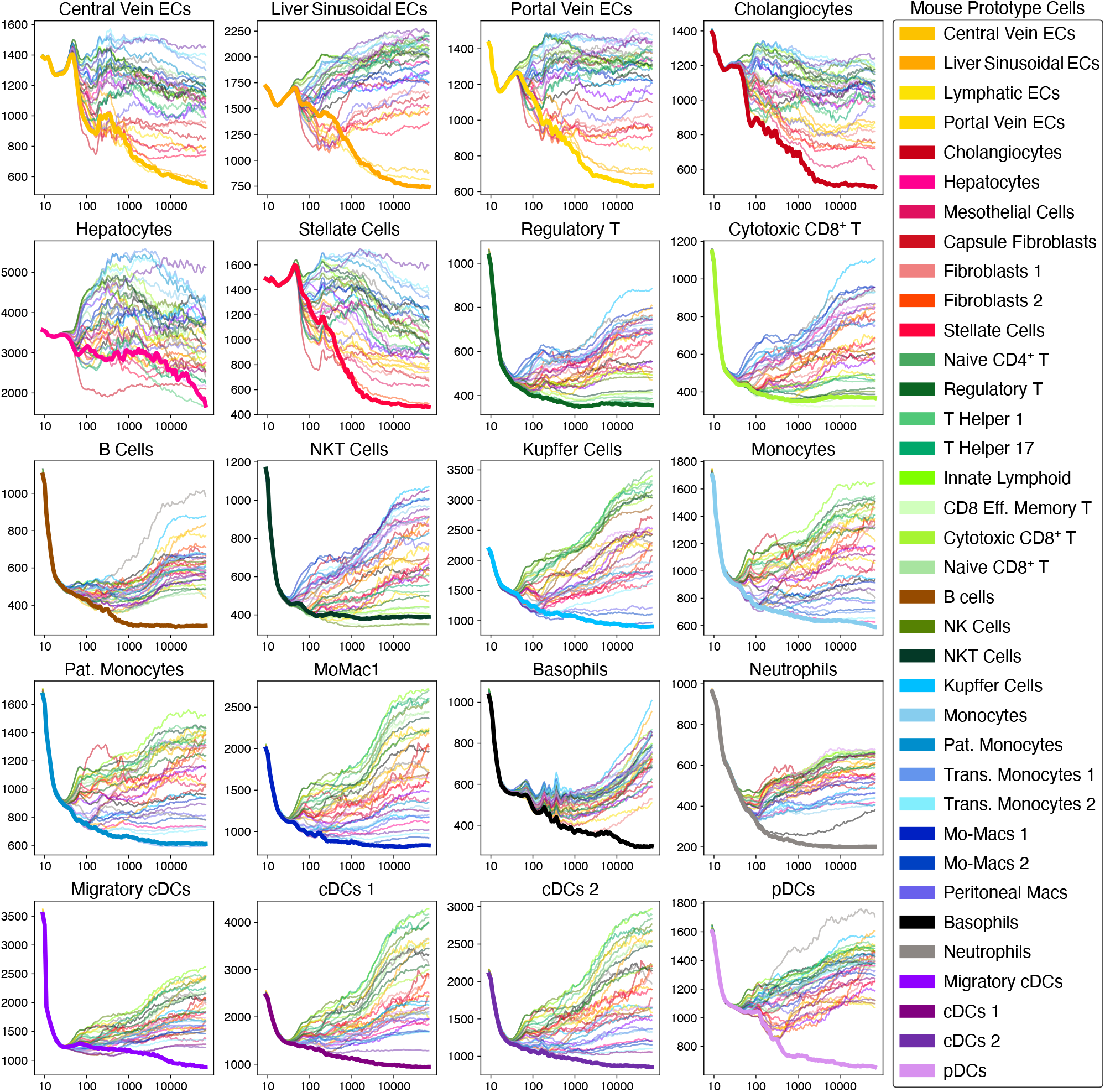
Illustration of the alignment process of scSpecies with *k* = 25 neighbors. On the y-axis, we plot the negative log-density values derived from reconstructing human liver cell prototypes using their candidate set of mouse latent variables. The x-axis shows a log-scale trajectory of these values, averaged over the last ⌈min(10, 0.05 × steps)⌉ iterations.

## Footnotes

1 Lower values indicate higher association.

